# Triangular Invariant Sets for Containment of Drug Resistance Under Evolutionary Therapy

**DOI:** 10.64898/2026.03.26.714636

**Authors:** Esteban A. Hernandez-Vargas

**Affiliations:** Department of Mathematics and Statistical Science, University of Idaho, Moscow, ID, USA

**Keywords:** biological systems, invariant-set analysis, population dynamics

## Abstract

Evolutionary therapies regulate heterogeneous populations by altering selective pressures through treatment sequences in cancer and infections. This letter develops an invariant-set framework for treatment-induced containment based on positive triangular invariant sets. For periodically switched systems, sufficient conditions are derived for the existence of such invariant regions. Robustness with respect to mutation is established by showing that the invariant simplex persists under small perturbations of the subsystem matrices. In the two-phenotype case, the analysis yields an explicit mutation threshold that separates regimes in which therapy cycling maintains containment from regimes in which mutation can enable evolutionary escape. Simulations illustrate the geometry of the invariant sets and the role of mutation and dwell time in containment robustness.

## I. Introduction

**A** Control-invariant set (CIS) is a region of the state space such that, for every initial condition in the set, there exists a control input that keeps the trajectory inside it for all future time [1]. This property makes control-invariant sets fundamental tools for the analysis and control of switched systems.

Switched systems are a class of hybrid systems in which a finite set of discrete modes selects the continuous dynamics governing the state evolution, together with a rule determining the switching among modes [2], [3]. The interaction between switching and continuous dynamics produces complex behaviors that complicate the characterization of invariant regions such as control-invariant sets (CIS).

The notion of periodic switching invariance introduced here is a direct specialization of classical positive invariance for discrete-time systems [1], [4], applied to the lifted system induced by a periodic switching signal. This problem has been investigated in several settings. Maximal control invariant sets for control linear switched systems were studied in [5], while nonlinear switched systems were considered in [6]. For autonomous linear switched systems, invariant sets have been analyzed both without dwell-time constraints [7], [8].

Over the last two decades, control-theoretic frameworks have been systematically applied to guide evolutionary dynamics under therapy switching [9]–[13]. In particular, control-invariant sets have been used to characterize the emergence of antibiotic resistance, providing a rigorous mathematical framework for analyzing how sequences of therapeutic interventions influence the containment of resistant populations [11], [12]. Collectively, these studies demonstrate that switched-system models, together with invariant-set concepts, provide a principled foundation for designing evolutionary therapies and steering population dynamics through adaptive treatment schedules [13].

Previous work has developed the concept of box invariance [14], which is a positively invariant region of the state space that contains the trajectories of the system. This problem remains decidable for important classes of nonlinear systems with polynomial structure. However, in the context of positive switched systems relevant to biological applications, this can be further simplified.

This letter introduces triangular invariant sets (see Fig. 1), which are particularly suitable for biological systems where the origin represents extinction of populations and must remain enclosed within the invariant region. Under this formulation, it is sufficient to define the coordinate axes corresponding to the system states that delimit the triangular region, significantly simplifying the construction of invariant sets. This geometric simplification facilitates the design of control invariant sets and provides a scalable framework for higher-dimensional positive systems. Conditions for switching and mutation robustness are derived, with explicit results for a two-phenotype model.

**Fig. 1:**
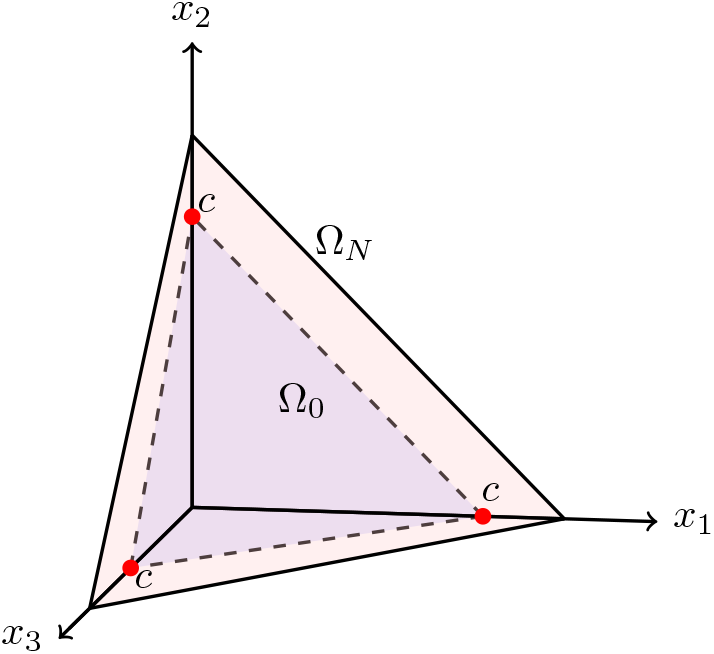
Triangular invariant set. The initial set Ω_0_ is formed by connecting the red points *c*, which can be contained in Ω_*N*_.

## II. PROBLEM STATEMENT

Biological populations under evolutionary therapy are naturally modeled in continuous time. To capture both therapy-dependent growth dynamics and phenotype transitions, the following switched positive system is employed

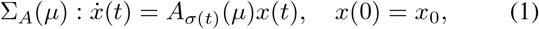

where

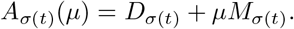

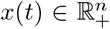 denotes the vector of phenotype populations, and *σ*(*t*) is the switching signal selecting the active therapy mode from the finite set ∑ := {1, 2, …, *q*}. The matrix *D*_*σ*(*t*)_ is diagonal and describes the therapy-dependent net growth or decay rates of each phenotype,

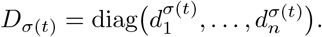

The matrix *M*_*σ*(*t*)_ ∈ {0, 1} ^*n*×*n*^ is a binary phenotype-transition matrix. An entry (*M*_*σ*(*t*)_)_*ij*_ = 1 indicates that phenotype *j* can transition into phenotype *i* under therapy *σ*(*t*), while (*M*_*σ*(*t*)_)_*ij*_ = 0 indicates that such a transition is not allowed. Typically, (*M*_*σ*(*t*)_)_*ii*_ = 0 since self-dynamics are already represented by the diagonal terms in *D*_*σ*(*t*)_.

The scalar parameter *μ* ≥ 0 quantifies the strength of phenotype transitions induced by mutation or phenotypic switching. When *μ* = 0, the system reduces to a purely diagonal switching system in which phenotypes evolve independently.

### Remark 1

*Since the matrices A*_*σ*(*t*)_(*μ*) *are Metzler for every mode σ*(*t*), *the switched system is positive: for any nonnegative initial condition* 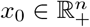, *the solution satisfies* 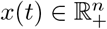 *for all t* ≥ 0 *[15]*.

Let *τ*_*k*_ := *t*_*k*+1_ − *t*_*k*_ denote the dwell time between switches. Defining 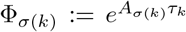, the system admits the discrete-time form

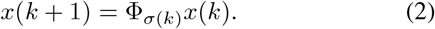

The following definitions are required before formally stating the problem.

### Definition 1 (Switching policy)

*A switching policy is a piecewise constant map σ* : ℝ_≥0_ → σ := {1, …, *q*} *satisfying σ*(*t*) = *σ*(*k*) *for t* ∈ [*kτ*, (*k* + 1)*τ*).

In many evolutionary therapeutic settings, treatments are applied in repeated cycles rather than arbitrary switching patterns. This motivates the following definition

### Definition 2 (Periodic switching)

*A switching signal is periodic if it repeats a fixed sequence of modes. For a cycle of modes* 1, …, *n, the system evolution over one period is given by the cycle map*

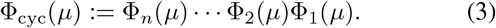

The following concepts are required to formalize invariance within this set.

### Definition 3

*(Invariant set [16]) A set* Ω ⊆ ℝ^*n*^ *is invariant for the system x*(*k* +1) = Φ(*x*(*k*)) *if x*(0) ∈ Ω *implies x*(*k*) ∈ Ω *for all k* ≥ 0.

### Definition 4 (Reachable set)

*The reachable set from x*_0_ *is the set of all states x*(*k*) *that can be reached from x*_0_ *under admissible switching signals σ*(*k*) *in a finite number of steps*.

### Definition 5

*(Controllable set [17]) A set C* ⊂ ℝ^*n*^ *is controllable if for any x*_1_, *x*_2_ ∈ *C there exists a switching sequence that steers the system from x*_1_ *to x*_2_ *in finitely many steps*.

Since the system is positive, points at *c* are placed on each coordinate axis and connected to form a simplex whose interior lies entirely within the positive orthant. Accordingly, the convex hull of the origin and these *n* axis points forms a triangular region. This construction is illustrated in three dimensions by the blue region in Fig. 1. The following definition formalizes the notion of a triangular invariant set.

### Definition 6 (Positive triangular invariant set)

*Let* Ω_0_ *be a subset* 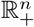 *defined as*

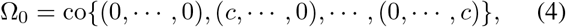

*where c >* 0. *If* Ω_0_ *satisfies Definition 3, then it is called a triangular invariant set. Equivalently, it can be written as*

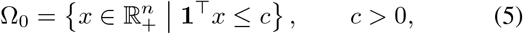

*where* **1** ∈ℝ^*n*^ *denotes the column vector of ones. The set* Ω_0_ *is therefore an n-simplex in the positive orthant. The symmetric construction arises from using the same parameter c along each axis, although symmetry is not required in general*.

### Main Contributions

Developing a triangular invariant set as in Fig. 1 provides a containment region for evolutionary therapy, ensuring that phenotype populations remain confined within a biologically meaningful positive domain under therapy switching. The main contributions of this study are:

- An invariant-set framework is formulated based on positive triangular sets in Definition 6. The resulting simplex geometry provides a simpler geometric alternative to box-based invariant constructions [18]. This reduction preserves the containment structure while enabling a more explicit and computationally tractable characterization of therapy-dependent admissible regions.
- Sufficient conditions under periodic switching are derived for triangular containment regions to persist in the presence of mutation. In the two-phenotype case, this yields an explicit mutation threshold that separates therapy regimes in which switching maintains evolutionary containment from regimes in which resistant populations can escape [10].

## III. Characterizing Triangle Invariant Sets

To characterize triangular invariant sets for the switched system (2), the backward-reachability construction algorithm proposed in [4] is adopted. Starting from the candidate simplex Ω_0_ defined in (5), the algorithm 1 iteratively computes the set of states that can be driven back to Ω_0_ under an appropriate switching choice. This procedure provides a constructive way to determine invariant and controllable regions for switched systems. For each subsystem map Φ_*i*_, define the inverse image of a set Ω ⊆ ℝ^*n*^ by 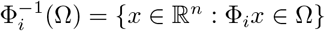.

### Algorithm 1

Backward construction of invariant sets [4].

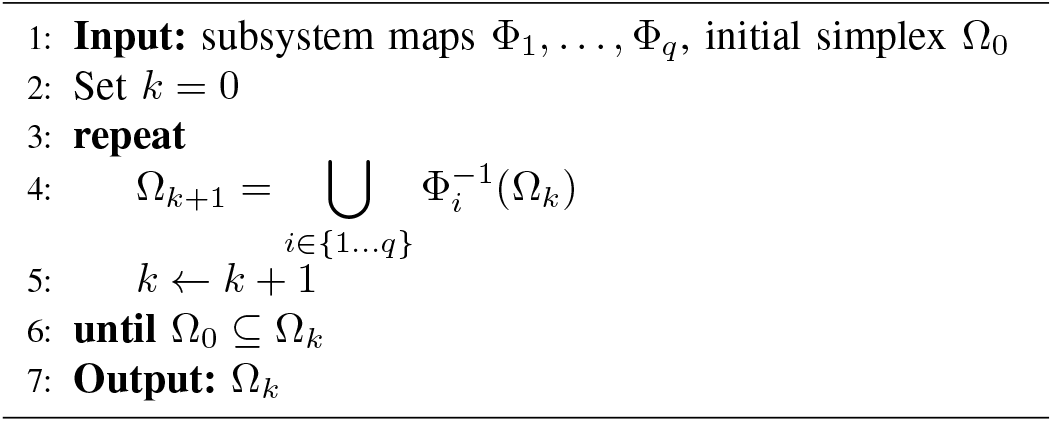

The set Ω_*k*+1_ consists of all states that can be driven into Ω_*k*_ in one step by at least one admissible mode. More generally, Ω_*k*_ represents the set of states that can be steered into the target set Ω_0_ within at most *k* switching steps. Therefore, the sequence {Ω_*k*_} forms a family of backward reachable sets associated with the triangular target region.

### Proposition 1

*If Algorithm 1 terminates in a finite number of steps, then the final set* Ω_*N*_ *is a controllable set for the switched system* (2). *In particular, for every x*_0_ ∈ Ω_*N*_ *there exists a switching sequence that steers the trajectory into the triangular target set* Ω_0_ *in finitely many steps*.

### A. Periodic Characterization of the Containment Region

The backward construction in Algorithm 1 provides a general procedure to compute the basin of containment, but it may generate unions of polyhedral preimages that are difficult to characterize analytically. For the class of positive switched systems considered here, the geometry can be described more explicitly by exploiting the periodic transition map induced by therapy cycling.

Under periodic switching (3), the dynamics over one cycle can be represented by a single transition map. Although this characterization provides a reduction of the full basin structure obtained from the backward construction, it enables explicit analytical conditions for containment. In particular, when the subsystem maps are diagonal, the simplex structure is preserved, and closed-form conditions for triangular invariance can be derived.

#### Theorem 2 (Diagonal Case)

*Consider the switched system* (1), *where each subsystem matrix* Φ_*i*_ ∈ ℝ^*n*×*n*^ *is diagonal. Assume that, for each switching mode i, the corresponding exponential matrix* Φ_*i*_ *has the form*

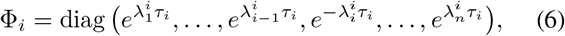

*where* 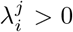 *for all i, j, and τ*_*i*_ *>* 0 *denotes the dwell time associated with mode i. Thus, in each mode, one diagonal entry (corresponding to index i) lies strictly inside the unit circle, while all others may exceed one. Let the candidate invariant set* Ω_0_ *be defined as in* (4). *Then, after one complete periodic switching cycle, the set satisfies* Ω_0_ ⊆ co(Ω_1_), *provided that, for each k* ∈ {1, …, *n*}, *the following condition on the eigenvalues holds:*

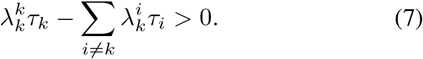

*Proof:* Assume *μ* = 0. Since each Φ_*i*_ is diagonal, the one-cycle map inverse is also diagonal and can be written as

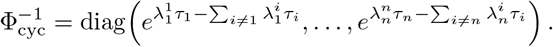

Define 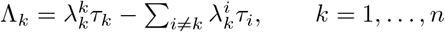.

If 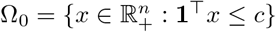, then its preimage under one cycle is

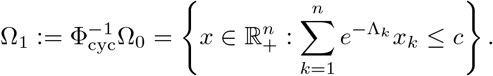

If Λ_*k*_ *>* 0 for every *k*, then 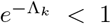 for every *k*, and therefore Ω_0_ ⊆ Ω_1_ ⊆ co(Ω_1_). This proves the claim.

#### Corollary 1

*Assuming constant dwell time between the switches τ* = *τ*_*k*_, *then, after one complete periodic switching cycle, the set satisfies* Ω_0_ ⊆ co(Ω_1_), *provided that, for each k* ∈ {1, …, *n*}, *the condition holds* 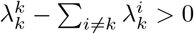.

*Proof:* It is straightforward considering *τ* = *τ*_*k*_ in (7).

Biological populations rarely evolve independently across phenotypes as assumed in Theorem 2. During therapy, evolutionary processes such as mutation, phenotypic switching, or horizontal gene transfer generate new variants and redistribute individuals across phenotypic compartments, thereby introducing coupling between population states and promoting adaptive resistance. In the model, this effect is represented by a small parameter *μ >* 0 that introduces off-diagonal couplings in the subsystem matrices of (1).

This raises the question of whether the invariant set obtained in the diagonal case persists under such perturbations. The following lemma shows that if the diagonal cycle maps Ω_1_ strictly inside itself, then this invariant geometry remains valid for sufficiently small perturbations of the system matrices.

#### Lemma 1 (Robustness under small perturbations)

*Let* Φ_cyc_(*μ*) *denote the state-transition map over one full switching cycle, as defined in* (3). *Assume that, for μ* = 0, *there exists a compact set* Ω_1_ *such that*

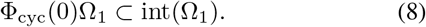

*If* Φ_cyc_(*μ*) *depends continuously on μ, then there exists* 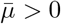*such that*

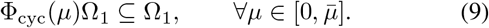

*Hence* Ω_1_ *remains invariant under the one-cycle map for all sufficiently small perturbations*.

*Proof:* Let *∂*Ω_1_ denote the boundary of Ω_1_ and define

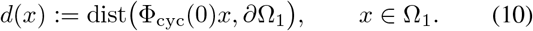

From (8), it follows that Φ_cyc_(0)*x* ∈int(Ω_1_) for all *x* ∈ Ω_1_, and therefore *d*(*x*) *>* 0 for all *x* ∈ Ω_1_. Since Φ_cyc_(0) is continuous and the distance to a closed set is continuous, the function *d* is continuous on Ω_1_.

Because Ω_1_ is compact, the Extreme Value Theorem implies that *d* attains its minimum on Ω_1_. Hence

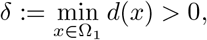

which means that the image Φ_cyc_(0)Ω_1_ stays uniformly away from the boundary of Ω_1_. Since Φ_cyc_(*μ*) depends continuously on *μ*, the map (*μ, x*) ⟼ Φ_cyc_(*μ*)*x* is continuous. Because Ω_1_ is compact, this continuity is uniform with respect to *x* Ω_1_. Therefore there exists 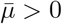 such that

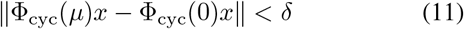

for all *x* ∈ Ω_1_ and 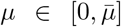. Let *x* ∈ Ω_1_ and set *y* = Φ_cyc_(0)*x*. Since dist(*y, ∂*Ω_1_) ≥ *δ*, the ball *B*(*y, δ*) is contained in Ω_1_. By (11), Φ_cyc_(*μ*)*x* ∈ *B*(*y, δ*), and therefore Φ_cyc_(*μ*)*x* ∈ Ω_1_. Thus 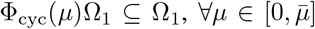, which proves (9).

#### Remark 2 (Evolutionary interpretation)

*The mutation parameter μ breaks the separability of the diagonal system by introducing weak coupling between phenotypes and enabling evolutionary mixing across compartments. For sufficiently small μ, these effects remain perturbative, and the invariant simplex persists under the cycle map. The admissible range of μ therefore quantifies the robustness of the triangular containment region. In evolutionary terms, the bound* 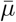 *represents a mutation–selection threshold: below this level, therapy switching maintains containment, whereas above it, mutation enables evolutionary escape*.

## IV. Mutation Threshold in Evolutionary Therapy: The Two-Phenotype Case

To illustrate how mutation perturbs the invariant-set structure, a two-dimensional model describing two interacting genotypes is considered. The dynamics switch between the two linear subsystems

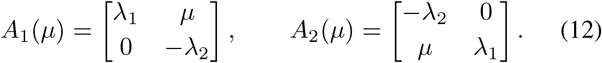

For the first mode, genotype 1 grows at rate *λ*_1_ while geno-type 2 declines at rate −*λ*_2_, with mutation coupling the two phenotypes. In the second mode, the roles are reversed, so genotype 2 grows while genotype 1 declines.

### Corollary 2 (Characterization of Two-Phenotype Model)

*Consider the two-mode switched system* (12) *with fixed dwell times τ*_1_, *τ*_2_ *>* 0. *Since A*_1_(*μ*) *and A*_2_(*μ*) *are triangular, their exponentials are*

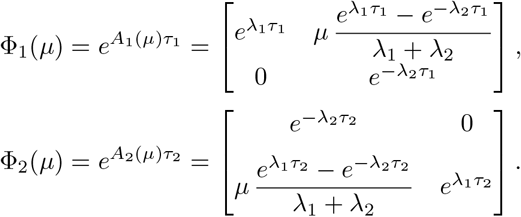

*Define the one-cycle map* Φ_cyc_(*μ*) := Φ_2_(*μ*)Φ_1_(*μ*).

*In the diagonal case μ* = 0, *let*

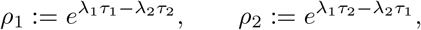

*with ρ*_1_, *ρ*_2_ *<* 1. *Let*

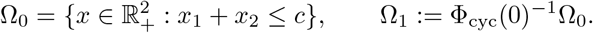

*Then*

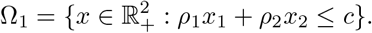

*Define*

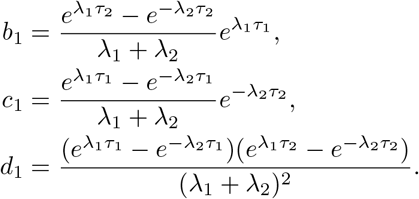

*Then* Ω_1_ *remains invariant for all mutation rates*

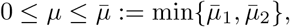

*where*

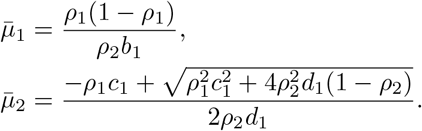

*Proof:* For *μ* = 0, Φ_cyc_(0) = diag(*ρ*_1_, *ρ*_2_), hence

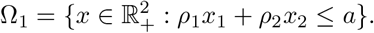

Using the expressions of Φ_1_(*μ*) and Φ_2_(*μ*),

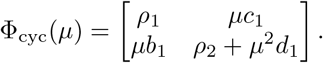

Since Φ_cyc_(*μ*) is positive, the simplex

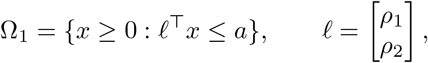

is invariant if and only if

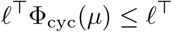

componentwise. This gives

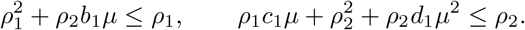

The first inequality is equivalent to

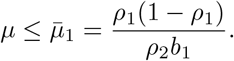

The second inequality is equivalent to

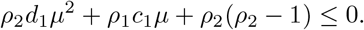

Since *ρ*_2_*d*_1_ *>* 0 and *ρ*_2_(*ρ*_2_ −1) *<* 0, this quadratic is precisely nonpositive for

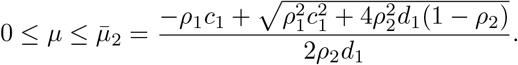

Therefore, for

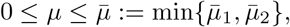

both inequalities hold, and thus Φ_cyc_(*μ*)Ω_1_ ⊆ Ω_1_.

### Remark 3 (Equal Dwell-Time Regime)

*In the equal dwell-time case τ*_1_ = *τ*_2_ = *τ, one has* 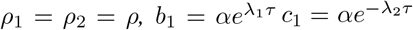, *and d*_1_ = *α*^2^, *where*

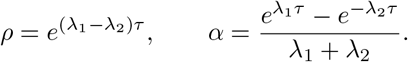

*Therefore Corollary 2 yields*

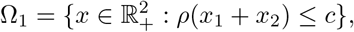

*and*

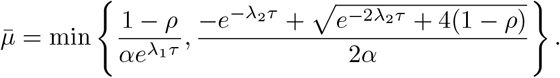

Thus, in the equal dwell-time case the invariant region becomes 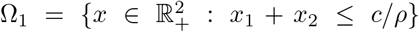, which is a scaled standard simplex. Geometrically, this means that the invariant constraint acts on the total population size rather than on the phenotype composition. In the context of alternating drug regimens, each treatment phase favors one phenotype while suppressing the other. Over a complete treatment cycle, the switching therapy therefore regulates the total population burden *x*_1_ + *x*_2_, effectively imposing a *treatment-induced containment region* on the system. Mutation redistributes population between phenotypes but, provided 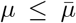, the total abundance remains confined within this containment region.

### A. Numerical Simulations

Numerical simulations are performed for the two-phenotype system (12) with parameters *λ*_1_ = 0.9, *λ*_2_ = 1.0, and *τ* = 1, with *c* = 1 defining the initial simplex Ω_0_ in (5). For these values, the admissible mutation bound is 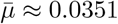.

#### Control Invariant Set

Algorithm 1 provides a stronger characterization of the containment region by iteratively computing backward reachable sets under each therapy mode. Starting from the initial simplex Ω_0_, successive iterations compute unions of preimages under the subsystem maps. From an evolutionary perspective, Ω_2_ in Fig. 2 defines a basin of evolutionary containment: the set of phenotype compositions for which therapy switching can still restore bounded population growth before mutation-driven escape occurs. Equivalently, Ω_2_ can be interpreted as a treatment rescue window, characterizing the population states that remain recoverable within two therapy phases.

**Fig. 2:**
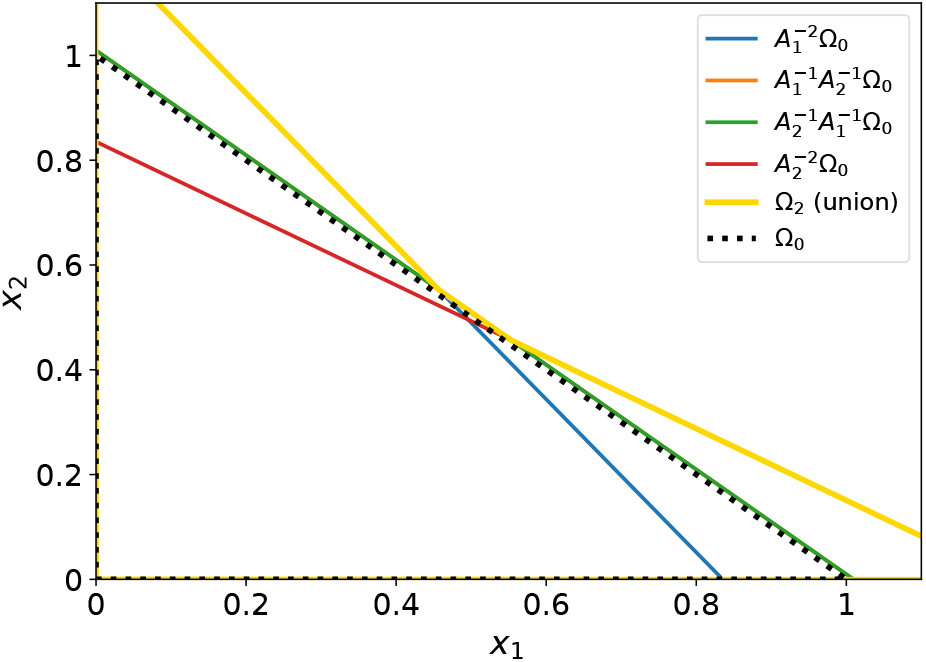
Invariant-set construction by the Algorithm 1.

#### One-cycle map

In contrast, the periodic characterization based on the cycle map yields a weaker but analytically tractable condition, enabling explicit analysis of the system and the derivation of mutation bounds.

In evolutionary terms, *μ* represents the mutation or phenotypic conversion rate between genotypes, while *λ*_1_ and *λ*_2_ encode their intrinsic growth and decay. The threshold 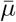 therefore acts as an *evolutionary robustness limit*: for 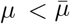, mutation introduces genetic variability without destroying the invariant population structure Ω_1_, corresponding to stable coexistence of types. As *μ* approaches 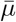, mutation pressure counteracts selection, the invariant geometry begins to deform, and beyond this point 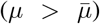 the population may lose bounded coexistence or shift dominance between genotypes.

This transition is illustrated in Fig. 3a. The blue triangle represents the invariant set Ω_1_ obtained from the diagonal case (*μ* = 0). The dashed orange curve shows the image of the boundary *∂*Ω_1_ under the cycle map for a mutation rate 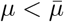, for which the image remains inside the simplex, preserving invariance. The dashed red curve corresponds to a mutation rate 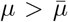, where part of the boundary image moves closer to the boundary of Ω_1_, illustrating the loss of guaranteed containment. This geometric comparison visualizes the mutation threshold 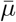 that ensures robustness of the containment region. Fig. 3b illustrates the phenotype dynamics for two mutation regimes. Solid curves correspond to a mutation rate below the theoretical threshold 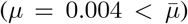, whereas dashed curves represent a larger mutation rate 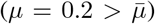. Blue curves denote phenotype *x*_1_ and orange curves denote phenotype *x*_2_. When the mutation rate is small, the evolutionary coupling between phenotypes remains weak and trajectories stay confined within the containment region predicted by the invariant simplex. In contrast, larger mutation rates strengthen the coupling between phenotypes and promote evolutionary escape from the containment regime under the switching therapy.

**Fig. 3:**
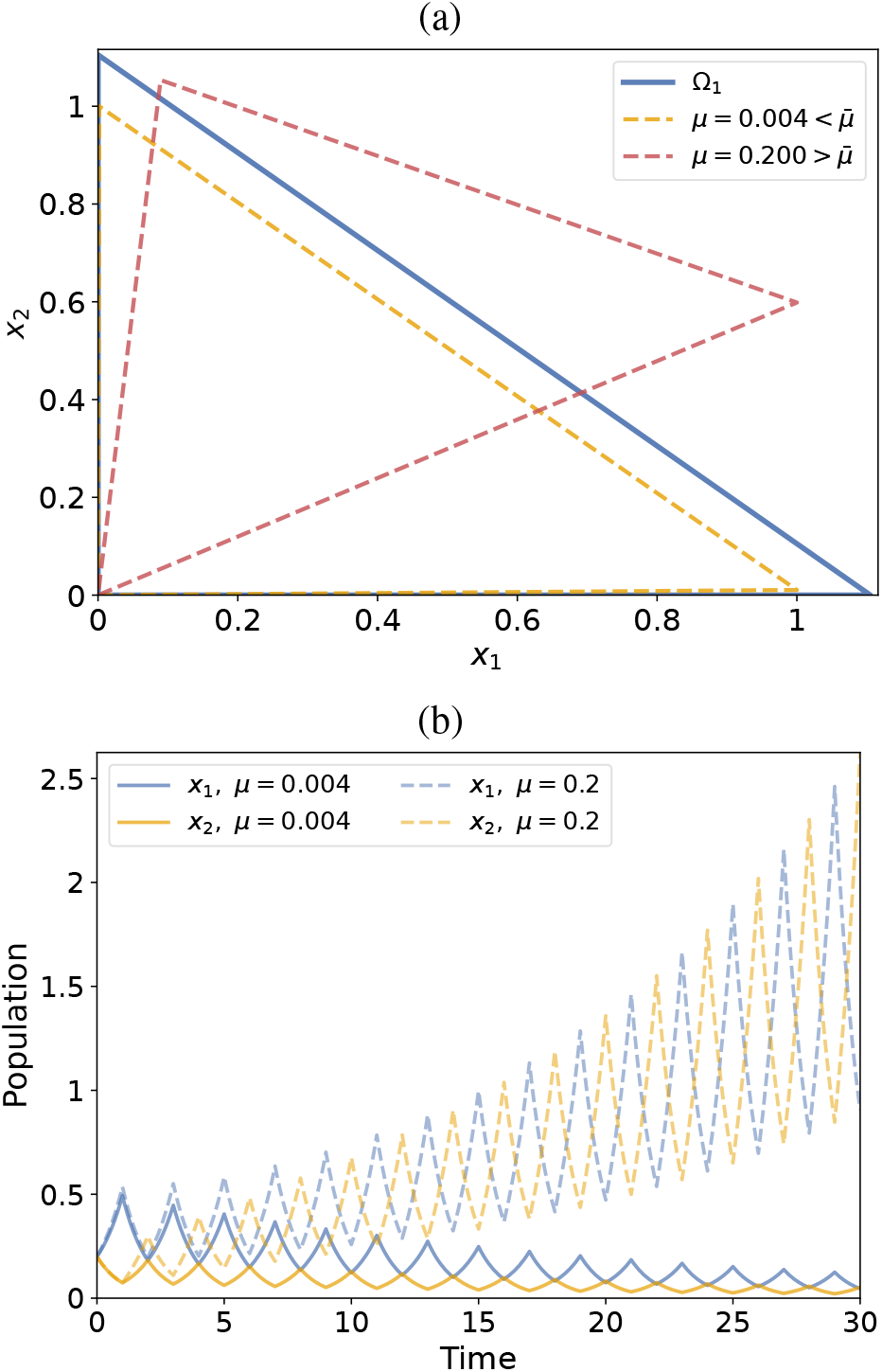
Panel (a) presents a one-cycle image of the containment simplex Ω_1_ under the perturbed cycle map Φ_cyc_(*μ*). Panel (b) shows the time evolution of the two phenotypes under periodic switching for two mutation rates.

Fig. 4 shows the maximal mutation rate for which the containment simplex Ω_1_ remains invariant under the cycle map Φ_cyc_(*μ*). As the dwell time increases, the admissible mutation range decreases, indicating that longer treatment phases amplify mutation-induced coupling between phenotypes and make the containment region more sensitive to evolutionary perturbations. Biologically, this means that prolonged exposure to a single drug regimen allows mutations to act over a longer time window before the therapy is switched. As a consequence, maintaining a treatment-induced containment region through cycling becomes more fragile when treatment phases are long.

**Fig. 4:**
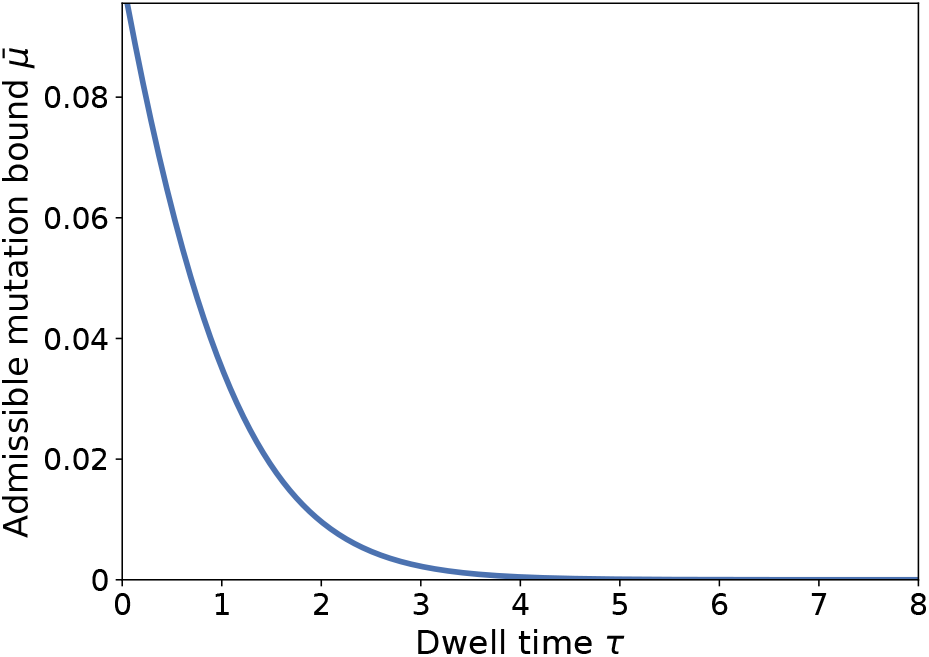
Admissible mutation bound 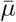 as a function of the dwell time *τ* in the equal dwell-time case *τ*_1_ = *τ*_2_ = *τ*.

## V. Conclusion

A triangular invariant-set framework was established for evolutionary therapy in positive switched systems. The proposed simplex-based containment region provides a biologically meaningful representation of bounded population burden under alternating treatment pressures, while preserving a tractable geometric structure. Under periodic therapy cycling, sufficient conditions were derived for containment in the absence of mutation, and robustness was shown to persist for sufficiently small phenotype-transition rates.

The numerical results illustrated how mutation and dwell time reshape the containment region and reduce the robustness of therapy cycling. These findings support the view that treatment schedules act not only through direct suppression, but also through their capacity to constrain evolutionary trajectories within admissible regions of phenotype space.

Future work will focus on adapting Algorithm 1 by [4] to exploit the geometric simplicity of triangular invariant sets, with the aim of obtaining faster and more scalable constructions of containment regions in higher-dimensional evolutionary systems.

